# Olfactory Co-Stimulation Influences Intranasal Somatosensory Perception

**DOI:** 10.1101/519439

**Authors:** Prasanna R. Karunanayaka, Jiaming Lu, Qing X. Yang, K. Sathian

**Affiliations:** Department of Radiology, The Pennsylvania State University College of Medicine, Hershey, PA, USA; Department of Neural and Behavioral Sciences, The Pennsylvania State University College of Medicine, Hershey, PA, USA; Department of Public Health Sciences, The Pennsylvania State University College of Medicine, Hershey, PA, USA; Drum Tower Hospital, Medical School of Nanjing University, Nanjing, China; Department of Neurosurgery, The Pennsylvania State University College of Medicine, Hershey, PA, USA; Department of Psychology, The Pennsylvania State University College of Medicine, Hershey, PA, USA; Department of Neurology, The Pennsylvania State University College of Medicine, Hershey, PA, USA

**Keywords:** multisensory integration, olfactory system, trigeminal system, localization

## Abstract

Olfactory sensitivity is influenced by intranasal trigeminal sensation. For instance, sniffing is central to how humans and animals perceive odorants. Here, we investigated the influence of olfactory co-stimulation on the perception of intranasal somatosensory stimulation. In this study, twenty-two healthy human subjects, with normal olfactory function, performed a localization task for weak air-puff stimuli, in the presence or absence of a pure odorant, phenyl ethyl alcohol (PEA; rose odor). Visual cues were used to inform participants to briefly hold their breath while weak, poorly localizable, air-puffs and/or PEA were delivered to either nostril. Although PEA alone could not be localized, when accompanied by a weak air-puff in the ipsilateral nostril, localization accuracy significantly improved, relative to presentation of the air-puff without the odorant. The enhancement of localization was absent when the air-puff and PEA were presented to opposite nostril. Since ipsilateral but not contralateral co-stimulation with PEA increased the accuracy of weak air-puff localization, the results argue against a non-specific alerting effect of PEA. These findings suggest an interaction between the olfactory and somatosensory trigeminal systems.

## INTRODUCTION

The intranasal trigeminal system processes chemosensory as well as somatosensory trigeminal stimulation conveyed via the trigeminal nerve. Although originating at receptors connected to fibers of the cranial nerve V (the trigeminal nerve) in the nasal mucosa, the two types of sensations seem to be relatively independent from each other. Olfactory information is processed via the olfactory nerve (cranial nerve I), but most odorants stimulate both sensory systems via mutually suppressing and enhancing interactions between each other (Brand 2006; Doty *et al.* 1978; Friedland and Harteneck 2017). For example, chemosensory trigeminal stimuli are perceived to be at higher intensities when presented with olfactory co-stimulation (Cain and Murphy 1980; Livermore *et al.* 1992). Further, evidence seems to support strong olfactory system interactions with the chemosensory trigeminal system; for instance, the threshold for chemosensory trigeminal perception is significantly impaired in subjects with anosmia (lack of odor perception) (Frasnelli *et* al. 2010; Frasnelli *et al.* 2007; Hummel *et al.* 1996; Hummel *et al.* 2003). Further, active sniffing as well as when odorants are delivered into participants’ nostrils using air-puffs (i.e., passive sniffing) improve olfactory perception (Frasnelli *et al.* 2009). Nevertheless, whether olfactory co-stimulation can modulate the intranasal somatosensory trigeminal perception still remains unclear.

One method to test the sensitivity of the trigeminal system is to perform a localization task in which a trigeminal stimulus is presented to one or other nostril and the subject is required to localize the stimulated nostril (Frasnelli *et al.* 2009; Hummel *et al.* 2003; Kleemann *et al.* 2009; Kobal *et al.* 1989). Correct localization indicates trigeminal perception, which is only possible if the trigeminal nerve is stimulated (Kobal *et al.* 1989). Experiments in this study used passive sniffing in which air-puffs were generated by blowing brief bursts of air into the nostril mimicking intranasal somatosensory stimulation. Air-puffs more or less resemble sniffing behavior which is an integral part of olfactory perception (Mainland and Sobel 2006; Sobel *et al.* 1998).

The goal of this paper is to investigate the influence of olfactory co-stimulation on intranasal somatosensory perception. Since multisensory integrative enhancement is most robust when unisensory inputs are weak, we designed our experiments following this principle of inverse effectiveness (Stein and Stanford 2008). For example, we used the pure olfactory odorant phenyl ethyl alcohol (PEA; rose odor) as the co-stimulant, which is not localizable during passive or active stimulation (Frasnelli *et al.* 2009; Kobal *et al.* 1989; Radil and Wysocki 1998). Since mixed olfactory/trigeminal stimuli are more easily localizable in the passive condition, we used weak air-puffs (Frasnelli *et al.* 2009; Porter *et al.* 2005; von Békésy 1964). Weak air-puffs are expected to evoke comparatively fewer neural impulses and therefore should be poorly localizable. If there is multisensory integration of olfactory and somatosensory trigeminal input. We expected substantial enhancement in air-puff localization accuracy under PEA co-stimulation, based on the principle of inverse effectiveness (Meredith and Stein 1986; Stein and Stanford 2008).

## METHODS

Twenty-two subjects (mean age 24.55 ± 3.36 years, 15 females) with normal smell function took part in the experiment after obtaining their written informed consent. The smell function was tested using the University of Pennsylvania Smell Identification Test (UPSIT) (Doty *et al.* 1984) and OLFACT: a test battery to evaluate odor identification and threshold. Unlike UPSIT, the OLFACT provides instructions, administers the stimuli, collects responses and scores them automatically. Study participants were recruited through advertisement in south-central Pennsylvania. The study had prior approval from the Pennsylvania State University College of Medicine Institutional Review Board and complies with the Declaration of Helsinki for Medical Research involving Human Subjects. Participants were also screened for complications that might lead to specific to olfactory dysfunction such as viral infection and allergies; such conditions led to exclusion from the study.

**Figure 1** shows the experimental setup, in which participants performed a two-alternative forced-choice (2AFC) discrimination to localize stimulation to the left or right nostril.

**Figure 1.**
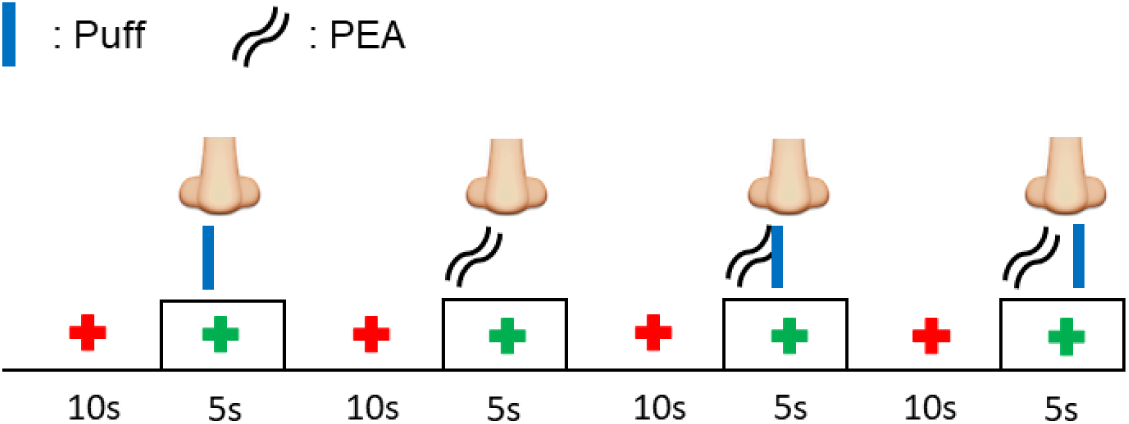
Nostril specific localization of weak air-puffs. A baseline air flow of 1 L/min was maintained throughout the experiment. Participants were instructed to focus on a ‘fixation cross’ that appeared on the screen. When the cross turned red (from white), they were instructed to hold their breath and attend to the stimulus. When the cross turned white again, subjects had to perform a left or right button press response to indicate the stimulated nostril. The experimental design took into account that “multisensory integration” requires simultaneous presentation of stimuli in order to enhance behavioral responses.

A detailed explanation is provided in the supplementary material (**Figure S1)**. In pilot studies, the intensity of air-puffs was varied by adjusting the duration of the valve open time (100-300 ms) and the peak air-flow of the puff (2-4 L/min) to find a weak intensity yielding a localization accuracy above chance but below the 75% correct level that is usually taken as threshold in a 2AFC; this intensity was used for the main experiment. The choice of intensity ensured that the stimuli were weak enough to be compatible with the inverse effectiveness principle, but not so weak as to be imperceptible. There were three experimental conditions in the main experiment. In one, weak air-puffs of a 100 ms duration were delivered randomly either to the nostril ipsilateral or contralateral to that receiving phenylethyl alcohol (PEA) stimulation. In a second condition, the weak air-puffs were delivered to either nostril without PEA. Third and fourth conditions comprised of PEA presented to either nostril. The inter-stimulus interval (ISI) was ∼15 sec. The total duration of the experiment was 10 mins. Visual cues were provided to inform participants to hold the breath in order to localize incoming weak air-puffs which were embedded in a constant flow of odorless-air and delivered bilaterally at a rate of 1 L/min.

Statistical analysis was performed using SPSS (IBM SPSS Statistics). First, we performed one-sample t-tests to determine if the localization score of each condition was greater than chance. Then, we used repeated-measures analysis of variance (ANOVA) to compare the results of localization during the various conditions, followed by paired t-tests for post hoc comparisons. We applied Bonferroni corrections for multiple comparisons, unless otherwise stated, and set the alpha value at 0.05.

## RESULTS

**Table 1** shows that the scores of olfactory function tests of the study participants were within the normal ranges for each test. Odor identification performance measured using the OLFACT and UPSIT was significantly correlated across participants (**Figure S2** of the supplementary material).

**Table 1.** Olfactory function characteristics of the study population

|  | OLFACT |  | UPSIT |
| --- | --- | --- | --- |
|  | Threshold | Identification | Identification |
| Mean ± SD | 11.47 ± 1.75 | 18.00 ± 1.45 | 36.23 ± 2.09 |
| Normal Range | $\geq 8$ | $\geq 16$ (Total 20) | $\geq 34$ (Total 40) |

We then analyzed whether participants were able to localize air-puffs in the different stimulation conditions. As shown in **Figure 2**, (for weak air-puffs alone), the accuracy of localization was 64.55% ± 3.48, which was significantly above chance (p < 0.001), but below the 75% correct threshold, as expected based on the choice of intensity from the pilot studies. When PEA alone was the stimulus, localization accuracy was 54.32% ± 3.07 which was not significantly above chance (p > 0.175), consistent with the prior literature indicating that pure odorant stimuli cannot be localized (Frasnelli *et al.* 2010). When the weak air-puff accompanied PEA in the ipsilateral nostril, localization accuracy was 83.64% ± 1.98, which was significantly above chance (p <0.001). The combination of the air-puff and the PEA in the contralateral nostril, yielded localization accuracy of 64.09% ± 3.90, again above chance (p < 0.001).

**Figure 2.**
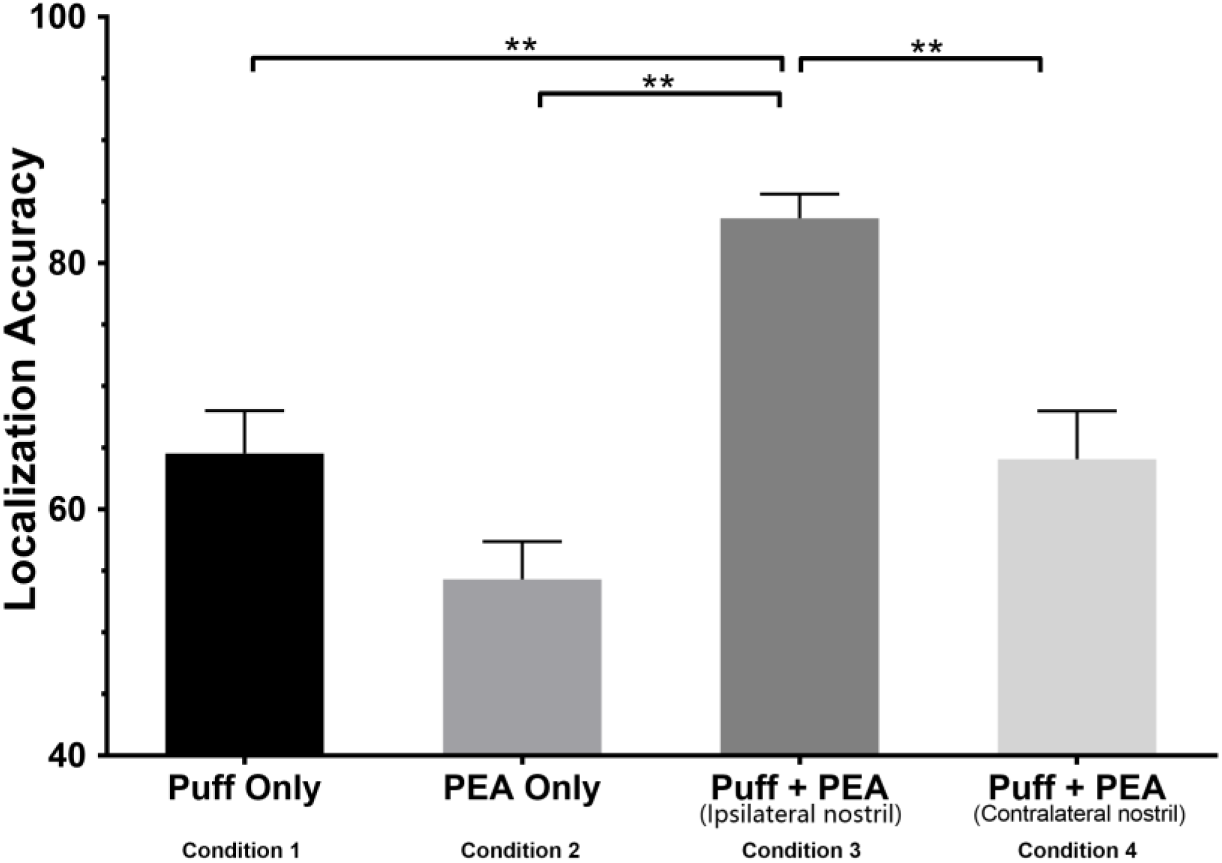
Nostril specific localization accuracy of weak air-puffs. Performance was significantly improved during co-stimulation with PEA in the ipsilateral nostril.

The ANOVA revealed a significant main effect of condition (F = 14.8, p < 0.001). Subsequent post hoc paired t-tests (Bonferroni-corrected α for six comparisons = 0.01) showed that the condition with PEA only differed significantly from other three conditions (p < 0.001). Additionally, ipsilateral PEA co-stimulation yielded significantly higher accuracy scores for weak air-puff localization than both the puff only and puff accompanied by PEA in the contralateral nostril condition (p < 0.01).

Finally, we investigated whether or not there are nostril specific differences during weak air-puff localization. As shown in **Figure 3**, no significant differences were found between the two nostrils in respective conditions.

**Figure 3.**
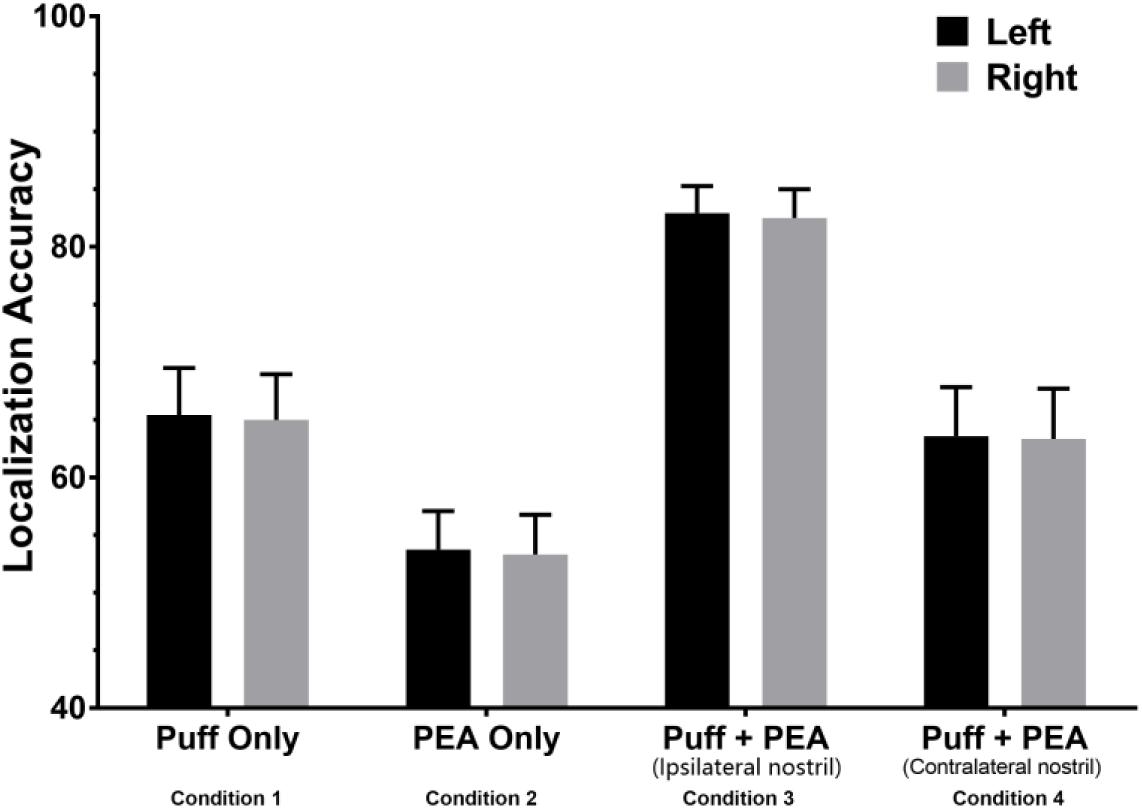
Left and right nostrils-specific localization accuracy of weak air-puffs showing no significant differences.

## DISCUSSION

Our results provide convincing behavioral evidence for the integration of olfactory and intranasal somatosensory information. We found that, although PEA (a pure odorant) could not be localized above chance, it enhanced the localizability of weak air-puffs delivered to the same, but not opposite nostril. Thus, the observed enhancement argues against a non-specific alerting effect by the odorant, which would have been expected to lead to similar effects whether PEA and air-puffs were delivered to the same or opposite nostrils. A similar enhancement has been reported using mixed olfactory and trigeminal (bimodal) stimuli (Tremblay and Frasnelli 2018). Together, these results are consistent with a general mechanism through which the olfactory system interacts with the trigeminal system, irrespective of whether it integrates chemosensory or somatosensory trigeminal information.

The mutual interactions between the trigeminal and olfactory systems seem to depend on several factors, including the quality and intensity of stimuli, and the concentration and duration of stimulation (Brand 2006; Hummel and Livermore 2002; Laing and Willcox 1987). Current results coupled with the findings of Tremblay et al. (2018) show that pure odorants can amplify the impact of somatosensory or chemosensory trigeminal stimuli on the trigeminal nerve, independent of the activated receptor type. Thus, the trigeminal system response seems to be independent of trigeminal receptors or type of stimulation (Kollndorfer *et al.* 2015). Therefore, measuring the sensitivity of one trigeminal stimulus may suffice as a general assessment of the trigeminal system (Frasnelli *et al.* 2011a).

Our study specifically focused on multisensory enhancement following the principle of inverse effectiveness, that is, enhancement is typically inversely related to the effectiveness of the individual cues that are being combined. This principle makes intuitive sense in this case and circumvents some of the criticisms of previous investigations. For instance, since strong air-puffs can be easily detected and localized, their combination with PEA would have a proportionately modest effect on neural activity and behavioral performance. In contrast, weak air-puffs would evoke comparatively fewer neural impulses and their responses would be subject to substantial enhancement when combined with PEA. In these cases, the multisensory neural responses from trigeminal as well as olfactory stimuli can exceed the arithmetic sum of their individual responses, as shown in the case of other stimulus combinations (Laurienti *et al.* 2005; Meredith and Stein 1986; Perrault *et al.* 2005; Stanford *et al.* 2005), enhancing the ability to localize the stimulated nostril. Therefore, future studies should appropriately choose stimuli when investigating olfactory-trigeminal interactions.

Ipsilateral, but not contralateral, olfactory co-stimulation with PEA increased the salience of air-puffs on the trigeminal system. This raises the possibility of olfactory and trigeminal interactions taking place in the nasal mucosa (Frasnelli *et al.* 2004), in addition to previously suggested central nervous system mechanisms (Cain and Murphy 1980). In the case of chemosensory trigeminal stimuli, Tremblay et al. (2018) suggested two candidate anatomical structures in which this interaction could take place: the mucosa of the nose and the olfactory bulb. The same structures may be relevant in the case of weak air-puff localization. This is because processing pathways for olfactory and air-puff stimuli are highly interconnected and share peripheral and central anatomical structures, despite activating different receptors and nerves at early processing sites.

Air-puff delivery is related to odor transduction which is an integral part of the process through which odorants bind to olfactory receptors (Mainland and Sobel 2006). Olfactory sampling or sniffing precedes this transduction stage. Indeed, air-puffs mimic sniffing behavior, which is necessary and sufficient for generating neural activity in olfactory brain regions such as the primary olfactory cortex (POC), a precursor to generating an olfactory percept of some sort, even in the absence of an odor (Mainland and Sobel 2006; Sobel *et al.* 1998). The source of the sniff-induced activation is the somatosensory stimulation that is induced by air flow through the nostrils (Mainland and Sobel 2006). Previous studies and our data show that somatosensory stimulation (or perception) is rapidly modulated in an odorant-dependent fashion. Therefore, given the anatomical structures involved in processing olfactory and trigeminal information, the mutual interactions between them are also likely supported by the central nervous system (Cain and Murphy 1980).

The lack of behavioral enhancement during contralateral PEA co-stimulation seems to contradict a predominant central nervous system interaction between the olfactory and trigeminal systems as suggested by Cain and Murphy (1980). One possibility is that weak air-puff localization score is not sensitive enough to central interactions. For instance, while simultaneous contralateral olfactory and trigeminal stimulation leads to the perception of stimuli with higher intensity, this may not translate into higher localization scores. Futures studies could confirm this hypothesis by looking at the correlation between weak air-puff intensity ratings and localization scores. Another possibility is that the site of the interaction between olfactory and trigeminal systems is dependent on the nature of the stimuli. In fact, previous studies have highlighted the fact that effects of co-stimulation are difficult to predict because they are dependent on the quality of stimuli (Hummel and Livermore 2002; Laing and Willcox 1987). Tremblay, et al., (2018), provides an enlightening discussion on this topic in relation to chemosensory trigeminal stimulation. They argue that the nature of stimuli (i.e., pure olfactory, mixed or pure trigeminal) plays a significant role and that future studies investigating olfactory-trigeminal interaction should choose stimuli appropriately.

The weak air-puff localization task did not show any inter-nostril differences. The intensity of air-puffs is likely the main stimulus parameter primarily responsible for localization. Nostril differences, such as favoring the right nostril, has been reported in odor discrimination (Savic and Berglund 2000; Zatorre and Jones-Gotman 1990; 1991), odor intensity judgement (Thuerauf *et* al. 2008) and odor localization (Frasnelli *et al.* 2009). Olfactory stimulation of the right nostril has also been shown to evoke higher activation in olfactory regions than stimulating the left nostril (Savic and Gulyas 2000), which is consistant with olfactory input being processed ipsilaterally (Hummel *et al.* 1995), at least prior to the POC. Nevertheless, nostril-specific behavior during chemosensory and somatosensory trigeminal stimulation remains unclear and needs to be thoroughly investigated in future research.

Most previous studies of olfactory-trigeminal integration used active sniffing during the localization task, where the odorant reaches the olfactory mucosa when participants take deep breaths. This technique precludes controlling for the intensity and duration of sniffs and therefore, the total amount of molecules reaching the nasal mucosa. In this case, participants may adapt their breathing patterns making the volume and/or vigor of inspiration unequal between conditions. Our study used passive stimulation where weak air-puffs and PEA were delivered within an air stream blown into the nostril, and the participants did not have to sniff in order for the stimuli to reach the olfactory mucosa. Since visual cues were provided to inform the participants to hold their breath in order to localize incoming weak air-puffs, we did not monitor their breathing patterns. Nevertheless, this is an important factor that needs to be controlled in studies of this nature as the trigeminal and olfactory systems work by integrating the total number of molecules over time (Cometto-Muniz and Cain 1998; Frasnelli *et al.* 2017; Frasnelli *et al.* 2011b; Wise *et al.* 2009).

In conclusion, using psychophysical techniques, we have shown that olfactory co-stimulation with a pure odorant does indeed influence processing of intranasal somatosensory stimuli. Ipsilateral, but not contralateral, co-stimulation increased the capacity to localize a somatosensory trigeminal stimulus. It remains for future work to establish the locus of this multisensory interaction and to clarify the underlying neural mechanism.

## ACKNOWLEDGEMENTS

The study was supported by the Leader Family Foundation, a grant from the U.S. National Institute of Aging (R01-AG027771) and the Department of Radiology, Penn State College of Medicine.

