## Supplementary Figures for "Olfactory Co-Stimulation Influences Intranasal Somatosensory Perception"

**SUPPLEMENTARY MATERIAL**

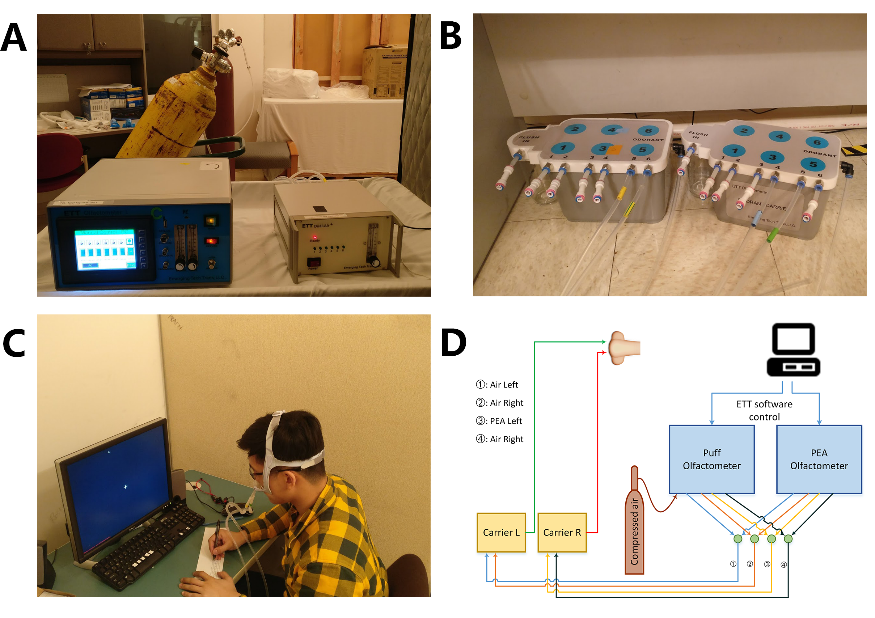

**Figure S1.** The experimental set up. (**A**): Olfactometers that delivered weak air-puffs, PEA and the baseline constant airflow of 1 L/min. Pilot studies were conducted to determine the intensity of air-puffs by (i) varying the duration of the valve open time between 100 to 300 ms and (ii) by varying the peak air-flow of the puff between 2 and 4 L/min; (**B**): Two odorant carriers were used to stimulate ipsilateral and contralateral nostrils; (**C**): E-prime visual presentation informing participants to hold the breath in order to localize incoming air-puffs embedded in the constant flow of odorless-air, delivered bilaterally at a rate of 1 L/min and (**D**): The overall experimental schematic.

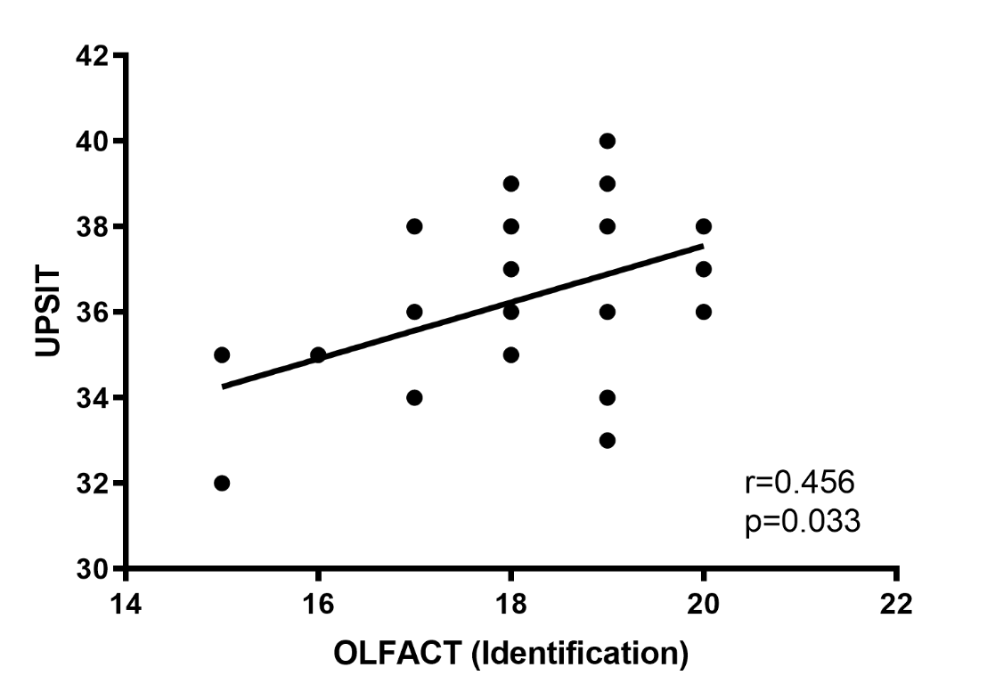

**Figure S2**. The correlation between UPSIT and OLFACT odor identification test scores.
